# Weight statistics controls dynamics in recurrent neural networks

**DOI:** 10.1101/475319

**Authors:** Patrick Krauss, Marc Schuster, Verena Dietrich, Achim Schilling, Holger Schulze, Claus Metzner

## Abstract

Recurrent neural networks are complex non-linear systems, capable of ongoing activity in the absence of driving inputs. The dynamical properties of these systems, in particular their long-time attractor states, are determined on the microscopic level by the connection strengths *w_ij_* between the individual neurons. However, little is known to which extent network dynamics is tunable on a more coarse-grained level by the *statistical* features of the weight matrix. In this work, we investigate the dynamical impact of three statistical parameters: *density* (the fraction of non-zero connections), *balance* (the ratio of excitatory to inhibitory connections), and *symmetry* (the fraction of neuron pairs with *w_ij_* = *w_ji_*). By computing a ‘phase diagram’ of network dynamics, we find that balance is the essential control parameter: Its gradual increase from negative to positive values drives the system from oscillatory behavior into a chaotic regime, and eventually into stationary fix points. Only directly at the border of the chaotic regime do the neural networks display rich but regular dynamics, thus enabling actual information processing. These results suggest that the brain, too, is fine-tuned to the ‘edge of chaos’ by assuring a proper balance between excitatory and inhibitory neural connections.

**Author summary:** Computations in the brain need to be both reproducible and sensitive to changing input from the environment. It has been shown that recurrent neural networks can meet these simultaneous requirements only in a particular dynamical regime, called the *edge of chaos* in non-linear systems theory. Here, we demonstrate that recurrent neural networks can be easily tuned to this critical regime of optimal information processing by assuring a proper ratio of excitatory and inhibitory connections between the neurons. This result is in line with several micro-anatomical studies of the cortex, which frequently confirm that the excitatory-inhibitory balance is strictly conserved in the cortex. Furthermore, it turns out that neural dynamics is largely independent from the total density of connections, a feature that explains how the brain remains functional during periods of growth or decay. Finally, we find that the existence of too many symmetric connections is detrimental for the above mentioned critical dynamical regime, but maybe in turn useful for pattern completion tasks.

## Introduction

In contrast to the artificial neural networks used in deep learning, which typically have a strict feed-forward structure, the networks of the brain contain many loops and are therefore recurrent in nature. This feature allows the cortex to maintain dynamical activity even without incoming external stimuli [1] and may therefore underlie such diverse operations as short-term memory [2–4], the modulation of neuronal excitability with attention [2, 5, 6], or the generation of spontaneous activity during sleep [7–9].

Recent micro-anatomical studies of the brain revealed that neural connectivity in the mammalian cortex has unique statistical properties. In particular, it was found that connections are sparse (low density), so that only a small fraction of possible connections are realized. The distribution of connection strengths is close to log-normal, and thus highly skewed, with a fat tail towards large magnitudes [10, 11]. Although the total number of non-zero connections can vary strongly between neurons, the ratio of excitatory to inhibitory connections is relatively constant [12]. Moreover, cortical networks contain a ‘skeleton’ of strongly connected neurons, linked pairwise in a bidirectional, symmetric way. This skeleton is embedded in a ‘sea’ of more weakly, non-symmetrically connected neurons [10].

Whereas the role of this peculiar connection structure is still purely understood, certain features seem to affect whether the brain can properly act as an information processor. For example, it has been shown that recurrent neural networks can show chaotic behavior for certain ratios between excitatory and inhibitory connections [1, 13]. It has even been speculated that certain social dysfunctions, such as autism and schizophrenia, are related to an elevated cortical excitation/inhibition balance [14]. Moreover, the discovered skeleton of neurons with strong bi-directional links may help to optimize information storage [15].

In a recent paper [16], we have investigated the relation between connectivity and system dynamics in small motifs of probabilistic neurons with binary outputs, assuming discrete, ternary connection strengths. We found that the balance between excitatory and inhibitory connections has a strong effect on the transition probabilities between successive motif states, whereas the total density of non-zero connections is less important.

Here, we extent our study to larger recurrent networks that consist of deterministic neurons with continuous outputs. Connection strength follow a random, log-normal weight distribution, but have prescribed values of the three control parameters density, balance, and symmetry. We analyze how these parameters affect the dynamical properties of the networks, in particular the Lyapunov exponent of the system trajectory in state space, the period length of cyclic attractors, and the cross correlation between individual neuron states.

As has been previously shown by Hopfield [19], networks with a very large fraction of symmetric bidirectional connections (symmetry parameter close to one) tend to end up in stationary fix points. We therefore focus on moderate and small symmetry parameters, and explore the two-dimensional phase diagram of system dynamics as a function of balance and density.

We find that this two-dimensional phase plane consists of three basic regions, corresponding to the possible attractors in deterministic and autonomous dynamical systems: periodic state cycles, chaos, and stationary fix point behavior. Strikingly, it is almost exclusively the balance parameter that controls in which of these three regimes a neural network is located, while the overall density of connections has a much weaker influence. In particular, the networks behave in a way that is suitable for information processing purposes only in a narrow range of balance parameters, located at the edge of the chaotic phase. This theoretical result is in line with the experimental finding that neural networks in the mammalian cortex have moderate degrees of symmetry and are tuned to rather specific values of balance, whereas connection density can vary widely between neurons and over time.

## Methods

### Neural network model

Our neural networks are based on simple deterministic neurons with zero bias (zero threshold). The total input *z_i_*(*t*) of neuron *i* at time *t* is calculated as:

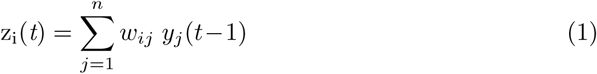

where *y_j_*(*t*−1) is the state of neuron *j* at time *t*−1 and *w_ij_* is the connection weight from neuron *j* to neuron *i*. The new state *y_i_*(*t*) of neuron *i* is computed as

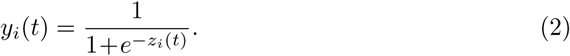

When simulating the dynamics of the networks, all neurons are updated simultaneously. The total state of a neural network at time step *t* can be summarized by the *n*-dimensional vector 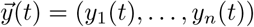, where *y_i_*(*t*) is the output state of neuron *i* at this time.

### Random weight matrix

The structure of a given neural network is defined by its weight matrix *W* = {*w_ij_*}. Here, we consider networks in which self-connections are forbidden, so that *w_ii_* = 0. For all non-zero matrix elements, the magnitudes of the weights are distributed according to a log-normal distribution,

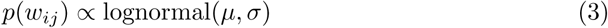

with location *µ* and scale *σ*.

### Statistical control parameters 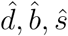

For a network with *n* neurons, the weight matrix has dimensions *n* × *n*. Due to the excluded self-connections, the diagonal elements of this matrix are zero, leaving a maximum possible number *n*(*n*−1) of non-zero matrix elements. We denote the actual number of non-zero weights by *m* = *m*_+_ + *m*_−_, where *m*_+_ and *m*_−_ are the numbers of positive and negative weights, respectively. Furthermore, we denote the number of non-zero matrix elements *w_ij_* for which a symmetric reverse connection *w_ji_* = *w_ij_* exists by *m_s_*. Based on these numbers, we define the *density* parameter 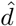, which varies between 0 for an unconnected and 1 for a fully connected network, by

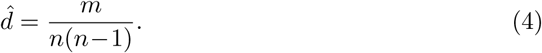

The *balance* parameter 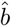, which varies between −1 for a purely inhibitory and +1 for a purely excitatory connection matrix, is defined by

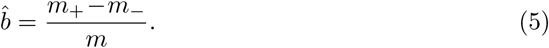

The *symmetry* parameter 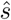, which varies between 0 for a completely non-symmetric and +1 for a completely symmetric (Hopfield-like [19]) network, is defined by

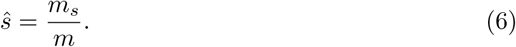

The meaning of these three control parameters is visualized in Fig.1.

**Fig 1.**
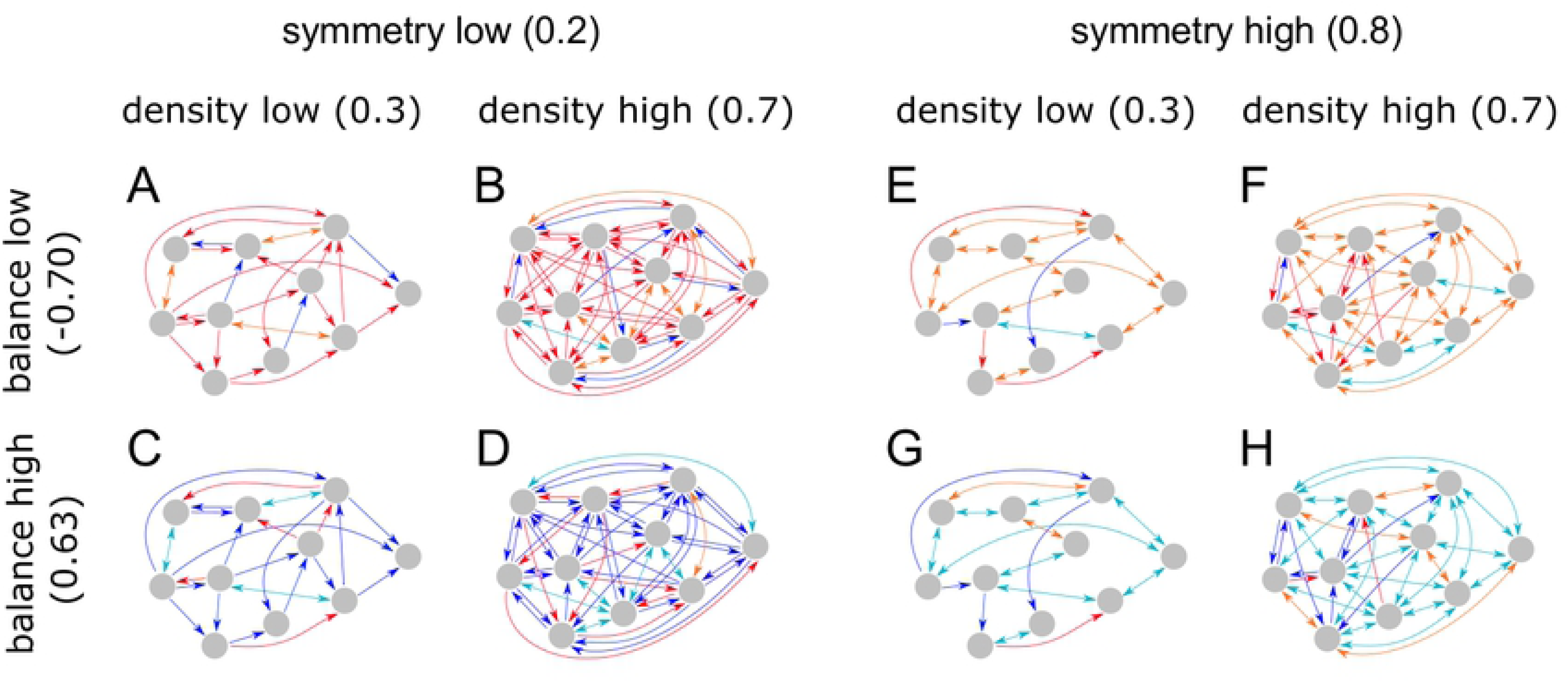
Visualization of the control parameters density 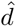, balance 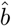, and symmetry 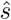 for eight example networks (A-H). Neurons are represented by gray circles, non-zero connections between neurons by arrows. One-headed arrows stand for uni-directional, two-headed arrows for bi-directional connections. Blue/magenta connections are excitatory (*w_ij_ >* 0), red/orange connections inhibitory (*w_ij_* < 0).

### Generation of weight matrices

Random weight matrices with prescribed values of the parameters 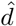, 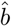, and 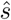 were generated in a series of steps. First, a fraction 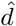 of the weights were drawn independently from a log-normal distribution with location *µ* = 0 and scale *σ* = 1, whereas all remaining weights were set to zero. Second, in order to introduce inhibitory connections to the network, a fraction 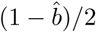 of the non-zero-weights were randomly selected and multiplied by −1. Third, the weights above the diagonal of the weight matrix were copied to below the diagonal, thereby creating a perfectly symmetric matrix. Finally, pairs of matrix elements below the diagonal were randomly selected and swapped iteratively, until the desired degree of symmetry 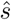 was achieved.

### Fraction of positive Lyapunov exponents *f*_λ>0_

Computing the new network state 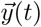 from the previous state 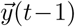 can be formally described by a vectorial update function

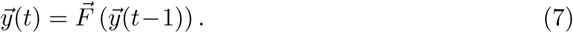

Starting from an infinitesimally close initial state 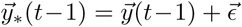 will lead to a different final state 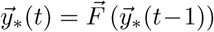. The sensitivity of the update function to this infinitesimal perturbation can be measured by the differential quotient

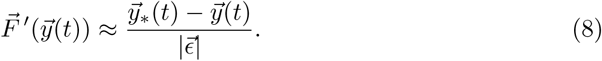

The *maximum Liapunov coefficient λ* of the update function is defined as

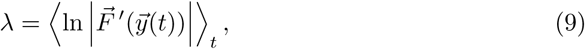

where 〈〉_*t*_ denotes the time average over all successive states of the system. It can be computed using well-established algorithms [17, 18]. A positive Lyapunov coefficient *λ >* 0 indicates that two nearby points in state space diverge exponentially, thus leading to irregular (chaotic) behavior. A zero or negative *λ* ≤ 0 indicates regular behavior. In general, within an ensemble of networks that are all characterized by the same set of control parameters 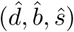, one finds *λ*-values of both signs. We therefore compute the *fraction of positive Lyapunov coefficients f*_λ>0_ for each ensemble and represent it as a color code in the ‘phase diagrams’ below.

### Average period length *T_av_*

Our recurrent networks are deterministic and autonomous dynamical systems. Thus, their trajectory 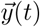 through *n*-dimensional state space is eventually governed by one of three possible attractors: a stationary fix point, a cycle of period *T*, or chaotic behavior. For each investigated network, we characterize the type of attractor by the measured period length *T*, that is, the number of time steps before the system state repeats itself for the first time 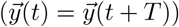. This includes the special cases *T* = 1, corresponding to a stationary fix point, and *T* = ∞, corresponding to a chaotic attractor. To identify repeating system states, we make use of a hash table. Since period lengths fluctuate for different networks from the same ensemble 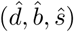, we compute the ensemble average *T*_av_ and use this average for color coding the phase diagrams.

### Root mean square of cross correlations *ρ*_rms_

The Lyapunov coefficient *λ* and the period length *T* characterize the long-time behavior of the neural networks. Another property that is relevant for a network’s information processing ability is the degree of correlation between individual neuron states *y_i_*(*t*) at the same time step *t*. For each pair *i,j* of neurons, it can be quantified by the Pearson cross correlation coefficient, defined as

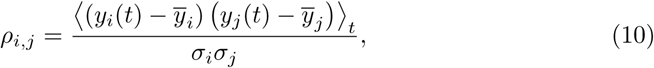

where 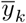 is the temporal mean of the time series *y_k_*(*t*) and *σ_k_* its standard deviation. In cases where *σ_i_* or *σ_j_* were zero, *ρ_i,j_* was set to 1. To characterize the global degree of correlation in a given neural network (without caring about the sign of the individual *ρ_ij_*), we computed the root mean square (RMS) over all neuron pairs

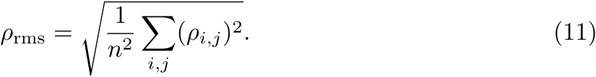

This quantity was additionally averaged over all members of a given 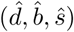 ensemble and then used for color coding the phase diagrams.

## Results

We first consider non-symmetric networks 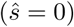, that is, networks without any bidirectional links of exactly the same strength. For each combination of balance 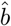 and density 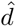 on a 11 × 10 regular grid, we generate an ensemble of 100 random networks. We then simulate the temporal dynamics of these networks, starting from random initial states. For each ensemble, we compute the fraction of positive Lyapunov coefficients *f_λ>_*_0_, the average period length *T*_av_, and the RMS of cross correlations *ρ*_rms_. The dependence of these dynamical quantities on the statistical control parameters is presented in the form of heat maps, which can be interpreted as dynamical ‘phase diagrams’ of these recurrent neural networks.

We initially focus on small networks of 100 neurons. When keeping the density close to one and gradually increasing the balance from negative to positive values, we find that the fraction of positive Liapunov coefficients *f*_*λ*>0_, indicating chaotic behavior, is close to zero, except for a narrow interval of balance values around 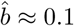. This chaotic interval broadens as the density parameter is reduced (Fig. 2A). In the 2D phase diagram, the chaotic regime therefore has an approximately triangular shape.

**Fig 2.**
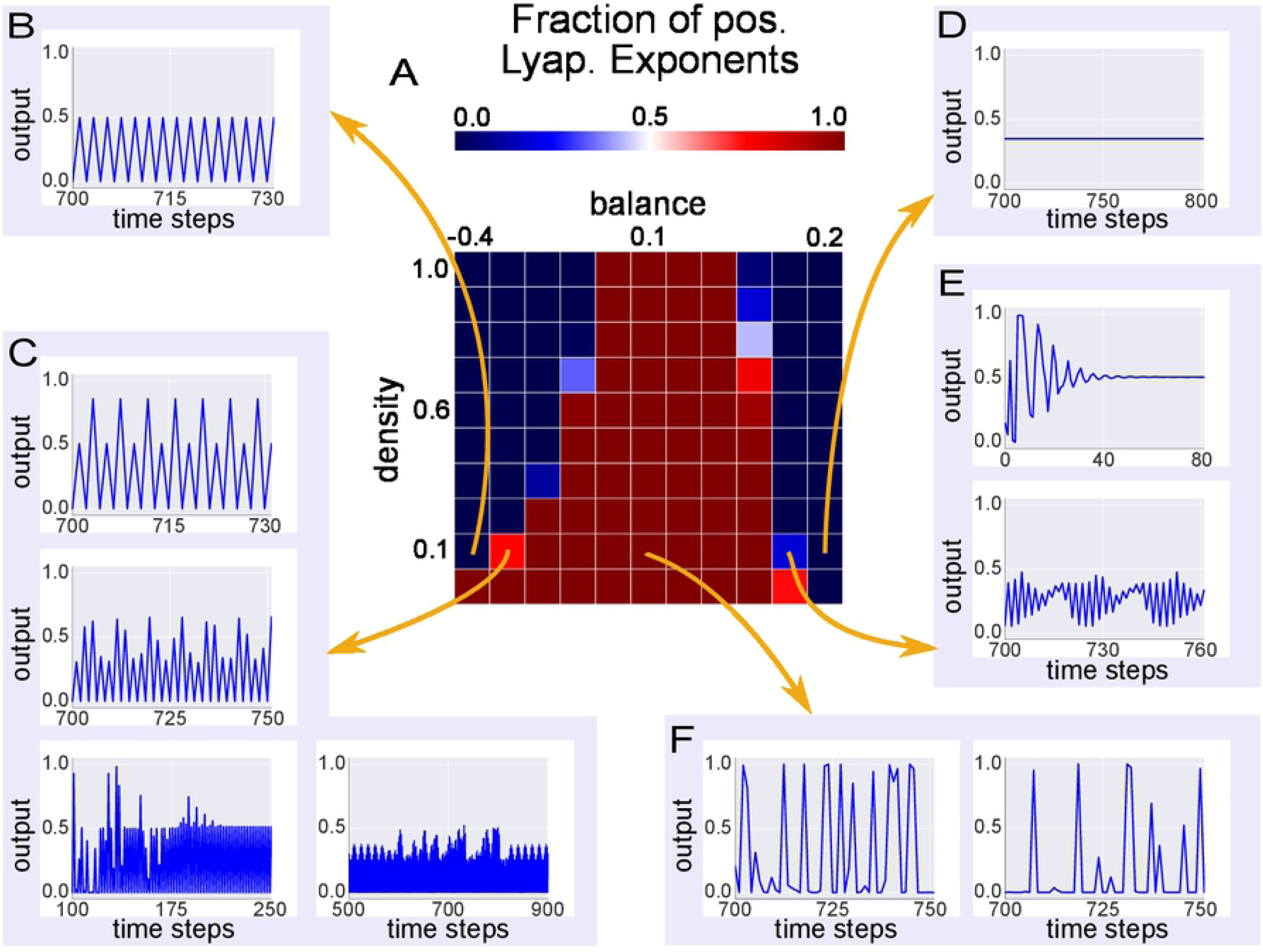
Dynamical phases in recurrent neural networks and characteristic output signals of individual neurons. (A) Two-dimensional phase diagram, showing the fraction of positive Lyapunov exponents *f_λ>0_* 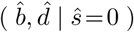 as a function of the control parameters balance and density, for a constant symmetry parameter 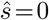. In the heat map, dark blue colors indicate *f_λ>_*_0_ ≈ 0, dark red colors *f_λ>_*_0_ ≈ 1. The red region in the center of the phase diagram is the chaotic regime, consistent with the irregular outputs of selected neurons (F). The ‘left’ blue region at negative balance values is the regime of cyclic attractors, often with small period lengths *T ≈* 2, as demonstrated with the neuron output (B). The ‘right’ blue region at positive balance values is the regime of fix points, as exemplified with the constant neuron output (D). The most interesting dynamics is found at the edge of the chaotic regime (C,E), where one finds cases of periodic behavior with large period length *T >* 2, periodic behavior with intermittent bursts, decaying oscillatory behavior, and ‘beating’ oscillatory behavior.

Inspecting the temporal output signals of selected neurons in the investigated networks (Figs. 2B-G), it turns out that the two regimes with *f_λ>_*_0_ ≈ 0 at the ‘left’ and ‘right’ side of the chaotic regime correspond to periodic attractors (Figs. 2B) and fix point attractors (Figs. 2D), respectively. The most interesting dynamics is found at the edge of the chaotic regime (Figs. 2C,E), where one finds cases of periodic behavior with large period length *T >* 2, periodic behavior with intermittent bursts, decaying oscillatory behavior, and ‘beating’ oscillatory behavior.

In a next step, we compare the phase distribution of *f_λ>_*_0_ with that of the other two dynamical quantities (middle and right column in Fig. 3). At the same time, we investigate the effect of system size (rows in Fig. 3, with different numbers of neurons *N*).

**Fig 3.**
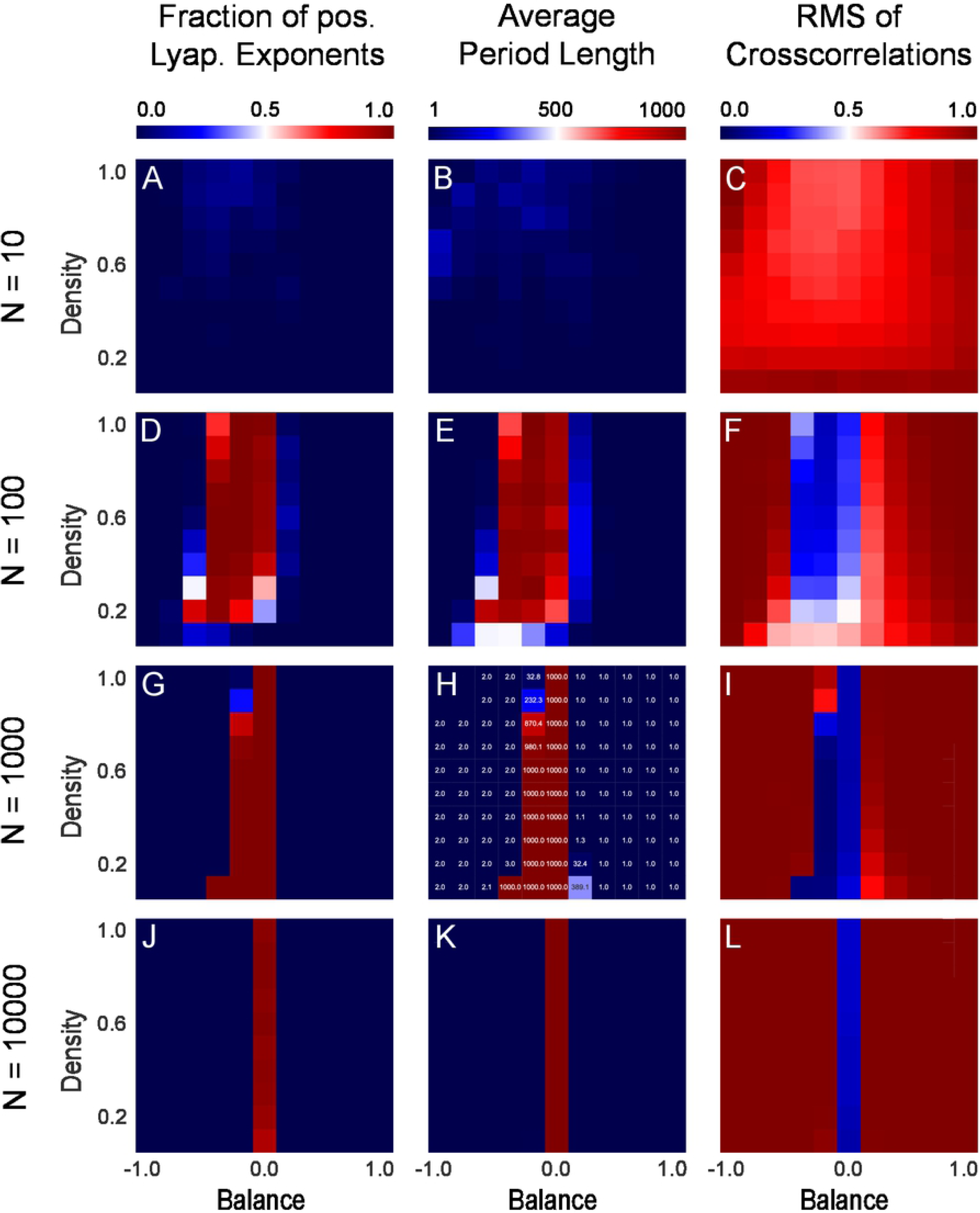
Comparing different dynamical measures, and the effect of system size. The columns correspond to the quantities *f_λ>_*_0_ (left), *T*_av_ (middle) and *ρ*_rms_ (right), as defined in the methods section. The rows from top to bottom correspond to increasing system sizes, characterized by the number of neurons *N* in the neural networks. For each of the 12 cases, a two-dimensional phase diagram is shown as a function of balance and density, keeping a constant symmetry parameter of 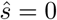. The three dynamic phases become apparent only for systems with a minimum size of *N* ≥ 100. The three different dynamical measures are mutually consistent. In particular, the chaotic regime is characterized by a *f_λ>_*_0_ close to one, by a diverging *T*_av_, and by a vanishing *ρ*_rms_. For large systems with *N* ≥ 10000, the density parameter has no more effect on the system dynamics, which is then controlled by the balance only.

We find that the three different dynamical quantities are mutually consistent. In particular, the chaotic regime is characterized by *f_λ>_*_0_ *≈* 1, by a diverging *T*_av_, and by a vanishing *ρ*_RMS_. The periodic regime is characterized by *T*_av_ *≈* 2 and by a relatively large *ρ*_rms_. The fix point regime is characterized by *T*_av_ = 1 and, again, by a relatively large *ρ*_rms_. Approaching the chaotic regime from either side by changing the balance parameter, *T*_av_ is rapidly increasing in the border region.

With increasing system size, the influence of the density parameter on the dynamical phase of the networks is diminishing. For large networks with *N ≥* 1000 neurons, the network dynamics is exclusively controlled by the balance parameter.

Finally, we investigate the effect of the symmetry parameter on the network dynamics (Fig. 4). By computing a complete 3D phase diagram of *f_λ>_*_0_ as a function of all three statistical control parameters, we find that balance and density have only an effect on the system dynamics when the symmetry is smaller than one, that is, when there are sufficiently many non-symmetric connections between the neurons. For a too large symmetry 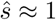, the system ends up in fix point attractors, irrespective of balance and density.

**Fig 4.**
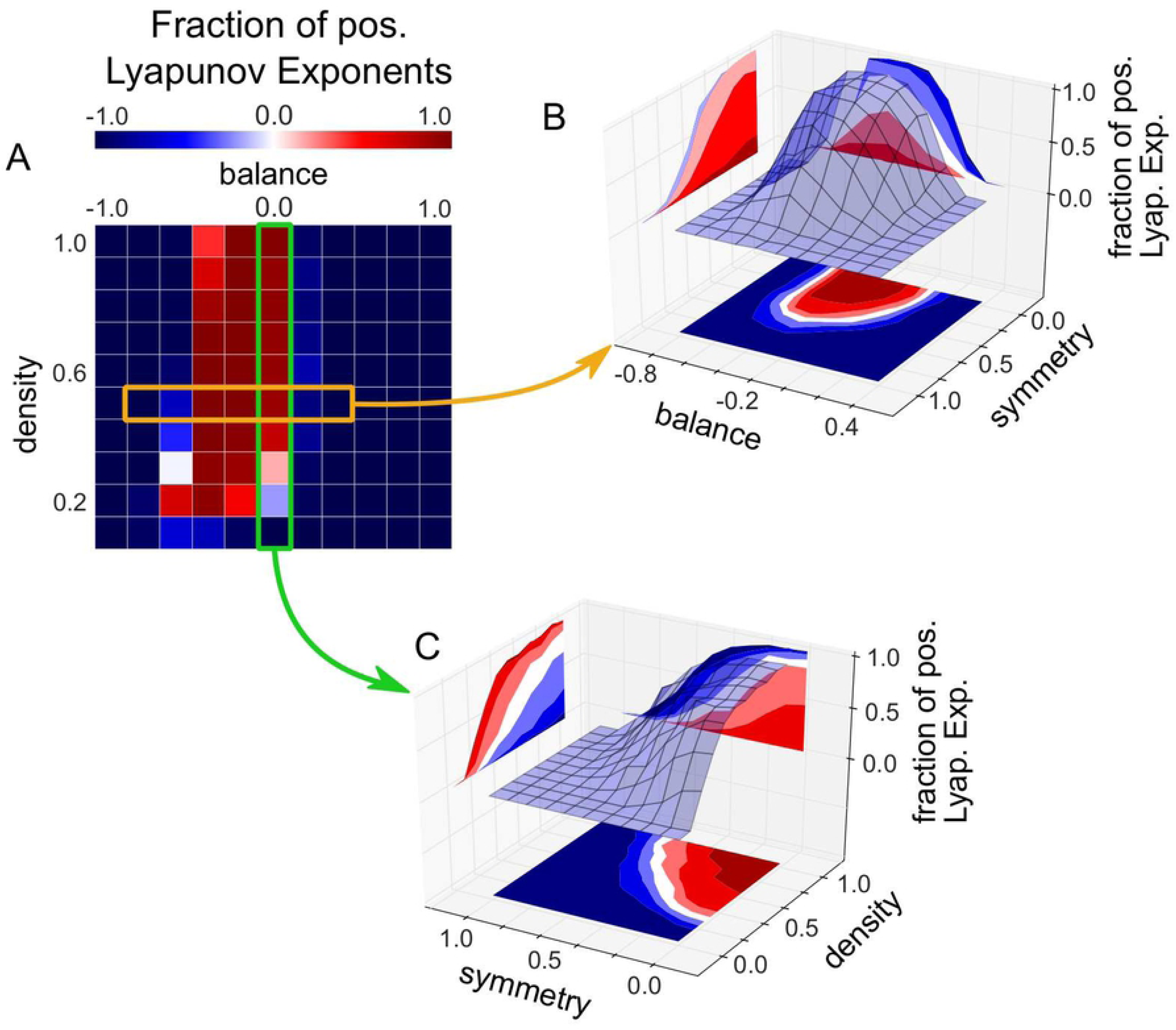
Effect of symmetry 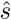 on system dynamics. (A): Standard plot of *f_λ>_*_0_ as a function of balance and density, for constant symmetry 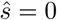. (B): Plot of *f_λ>_*_0_ as a function of balance and symmetry, for constant density 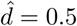 (see orange box in (A)). (B): Plot of *f_λ>_*_0_ as a function of symmetry and density, for constant balance 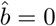 (see green box in (A)). For too large symmetry 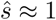, the system ends up in fix point attractors, irrespective of balance and density.

## Conclusion

In this work, we have demonstrated that the dynamical behaviour of recurrent neural networks can be effectively tuned by certain statistical properties of the network’s connection weight matrix.

In particular, a large fraction of symmetric, bi-directional neural connections 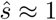 favors fix point attractors, and may therefore be useful for pattern completion tasks, as in the Hopfield model [19]. However, rich dynamical behavior is only possible for moderate or small degrees of symmetry.

For those non-symmetric networks, the statistical parameter with the largest impact on system dynamics is the balance 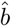 [16]. This ratio between excitatory and inhibitory connections controls, with high fidelity, whether a free-running neural network will behave stationary, oscillatory, or irregularly. Moreover, fine tuning of the balance parameter can bring the system to the edge of the chaotic regime, where the outputs of the neurons produce complex wave forms, and where the system may depend sensibly, but still regularly, on external inputs. We speculate that this regime is most suitable for purposes of neural information processing [20–24], and that biological brains may therefore control the parameter 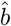 in a homeostatic way [1, 25, 26].

By contrast, the impact of the overall connection density 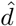 on network dynamics, at least in realistically large systems with many neurons, is much smaller than that of the balance 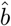. In principle, a recurrent neural network can gain or loose a large random fraction of neural connections without changing its dynamical attractor state, as long as the balance 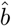 remains unchanged. This surprising robustness, for which the term graceful degradation has been coined [27], may help to keep the cortex functional during periods of growth and decay.

Future work will need to clarify how recurrent neural networks, statistically tuned into specific attractor states, react to external inputs. A particularly interesting question will be whether the edge of chaos is also marked by a large mutual information between input signals and the internal sequence of states within the recurrent neural network.

## Acknowledgments

This work was supported by the Deutsche Forschungsgemeinschaft (DFG, grant SCHU1272/12-1). The authors are grateful for the donation of two Titan Xp GPUs by the NVIDIA GPU Grant Program.

